# Generation of High-Affinity Anti-GIPR Antagonist Antibodies with Sustained and Non-rebound Weight Loss in DIO Mice by AlfaBodY

**DOI:** 10.64898/2026.04.21.719783

**Authors:** Yongkang Long, Zhangzhan Xu, Ning Zhang, Gaojian Chen, Weihai Chen, Zhongwei Chen, Ao Wang, Zhiming Liang, Yingya Wang, Yi Zeng, Kingsley Leung, Liang Chen

## Abstract

The glucose-dependent insulinotropic polypeptide receptor (GIPR) is an attractive therapeutic target for metabolic disorders, with GIPR antagonism emerging as a promising strategy for obesity and type 2 diabetes. However, developing functional antibodies against GPCRs remains challenging due to their complex architecture and conformational dynamics. Here, we employed AlfaBodY, an iterative active learning platform integrating structural and sequence information, to *in silico* design human anti-GIPR antibodies. Through four rounds of optimization, we generated antibodies with high binding affinities. Lead candidates AB106-131 (K_D_ 1.2 nM) and AB106-156 (K_D_ 1.7 nM) exhibited 7 to 10-fold higher affinity than 2G10 (K_D_ 12 nM) while maintaining comparable antagonistic activity in a cAMP reporter assay (IC_50_ 4∼5 nM). In diet-induced obese mice, AB106-156 alone induced weight loss comparable to that of semaglutide (∼ -15%), while preserving lean mass and achieving sustained weight control after treatment withdrawal. Co-administration with the GLP-1 receptor agonist semaglutide produced synergistic weight reduction (−25.4%) and markedly attenuated the fat-mass rebound observed with semaglutide alone. Our results demonstrate that AI-driven design can generate potent anti-GIPR antibodies with favourable *in vivo* efficacy, supporting further development of GIPR antagonist for obesity and related metabolic disorders. The AlfaBodY platform enables faster development of more efficacious biologic drugs.

## 1 Introduction

The glucose-dependent insulinotropic polypeptide receptor (GIPR) is a class B G protein-coupled receptor (GPCR) that plays a pivotal role in metabolic regulation^[1]^. GIPR mediates the physiological effects of glucose-dependent insulinotropic polypeptide (GIP), a key incretin hormone secreted by intestinal K cells in gut. Upon activation by its endogenous ligand, GIP, the receptor stimulates insulin secretion from pancreatic β-cells in a glucose-dependent manner, while also exerting pleiotropic effects on adipose tissue, bone metabolism, and the central nervous system^[2, 3]^. These physiological functions position GIPR as an attractive therapeutic target for metabolic disorders such as type 2 diabetes and obesity^[4-6]^. Pharmacologically, GIPR activation promotes glucose-dependent insulin secretion, supports pancreatic β-cell function, and regulates lipid metabolism in adipose tissue. Therefore, GIPR has garnered considerable attention as a key target in obesity pharmacotherapy, with an increasing number of candidates advancing from preclinical development into clinical-stage evaluation^[6]^ and market approval.

Another gut-derived incretin hormones are glucagon like peptide-1 (GLP-1), known for their ability to augment glucose-stimulated insulin secretion and to promote satiety. Over the past two decades, GLP-1R agonists (GLP-1RA) have evolved from short-acting native peptides requiring continuous infusion to long-acting, once-weekly formulations exemplified by semaglutide and dulaglutide, with recent approvals for chronic weight management (Wegovy HD® semaglutide 7.2 mg) demonstrating unprecedented efficacy, achieving mean weight reductions exceeding 18% at 68 weeks in STEP UP clinical trials ^[7]^. Notably, dual agonists that simultaneously activate GIPR and GLP-1R, such as tirzepatide, have demonstrated superior weight-loss efficacy compared with GLP-1R monoagonists, underscoring the therapeutic potential of integrated GIPR-based strategies in metabolic disease management. In the SURMOUNT-5 trial, tirzepatide demonstrated a mean weight reduction of -20.2% at week 72, compared to -13.7% for semaglutide^[8]^.The therapeutic concept for targeting GIPR has evolved considerably. While initial research focused on GIPR agonism to enhance insulin secretion, genetic studies demonstrating that individuals with loss-of-function GIPR variants exhibit favorable metabolic phenotypes—including reduced body weight and improved insulin sensitivity—have substantiated the therapeutic potential of GIPR antagonism^[9, 10]^. This concept gained clinical validation with the development of maridebart cafraglutide (AMG133, 2G10 mAb conjugated GLP-1RA). Unlike the dual-agonist approach, AMG133 is a first-in-class antibody-peptide conjugate (APC) that integrates GIPR antagonism with GLP-1 receptor agonism^[11, 12]^. Recent Phase 2 clinical data demonstrated superior weight loss efficacy of up to 20% over 52 weeks with a differentiated monthly dosing profile^[13]^ positioning GIPR antagonism as a potent challenger in next-generation incretin-based therapies. Compared with tirzepatide, a dual GIPR and GLP-1R agonist, AMG133 offers a longer dosing interval by utilizing an APC backbone, which may improve patient adherence. Therefore, enhancing the activity of GIPR antagonists to further augment their weight-loss efficacy would be of great significance.

Despite this progress, developing therapeutic antibodies against GPCRs remains challenging due to their complex transmembrane architecture, conformational flexibility, and the limited immunogenicity of their extracellular domains^[14, 15]^. The antigen generation itself is difficult, as producing correctly folded, conformationally relevant GPCR protein for immunization or screening requires advanced stabilization strategies^[15]^. Conventional discovery platforms—such as hybridoma technology or phage display—are often time-consuming and labor-intensive, limiting the rapid development of candidates against such structurally intricate targets^[16]^.

In recent years, computational and AI-driven platforms have enabled the *de novo* design of antibodies with tailored binding properties, offering a more efficient and targeted approach to therapeutic antibody development^[17]^. Recent advancements in computational biology and artificial intelligence have revolutionized the field of therapeutic antibody design. Platforms and tools like AlphaFold^[18, 19]^ for protein structure prediction, RFdiffusion^[20, 21]^ for protein design, and generative models such as BoltzGen^[22]^ enable the *de novo* design of antibodies with tailored properties, offering a more rational and efficient approach. In this study, we utilized our AlfaBodY platform—an iterative active learning system that integrates structural and sequence information to design novel anti-GIPR antibodies. This platform simultaneously optimizes both the complementarity-determining regions (CDRs) and framework regions, aiming to generate antibodies with high binding affinity favorable functional activity in cellular assays, and promising efficacy in preclinical animal models, its performance exceeds existing benchmark, such as the reference antibody of 2G10. Critically, our design process incorporated a developability assessment from the outset using AlfaDAX, our proprietary *in silico* platform. We aimed to mitigate common issues in therapeutic antibody development, such as aggregation propensity, high viscosity, and non-specific binding, by employing predictive algorithms.

A series of GIPR antagonist antibodies were generated and investigated *in vitro*, following systematic evaluation the lead candidate was confirmed and named as AB106-156. AB106-156 is a humanized IgG1 monoclonal antibody with high affinity to cynomolgus monkey and human GIPR but does not bind to mouse or rat GIPR, a *GIPR* humanized transgenic animals (B-*hGIPR* mice) were therefore employed to assess its effect in pharmacology studies in mice. The objectives of the current study were to characterize the antibody generation process, along with its *in vitro* activities. In addition, the synergistic effect resulting from the combination of GIPR antagonism and GLP-1R agonism was investigated, supporting the subsequent development of GIPR antagonist antibody and GLP-1 peptide conjugates.

## 2 Results

### 2.1 Design of anti-GIPR antibodies via AlfaBodY platform

Our AlfaBodY platform utilizes an iterative active learning process to *in silico* design antibody sequences by refining candidates over multiple cycles. This method integrates both structural data and sequence information to achieve high-affinity binding to a specific target. The process begins with a template antibody sequence and the target protein. This serves as the foundation for generating new antibody sequences. Unlike methods that focus only on specific regions, AlfaBodY is used to design both the CDR (Complementarity-Determining Region) and the Framework sequences of the antibody (Figure 1A).

**Figure 1.**
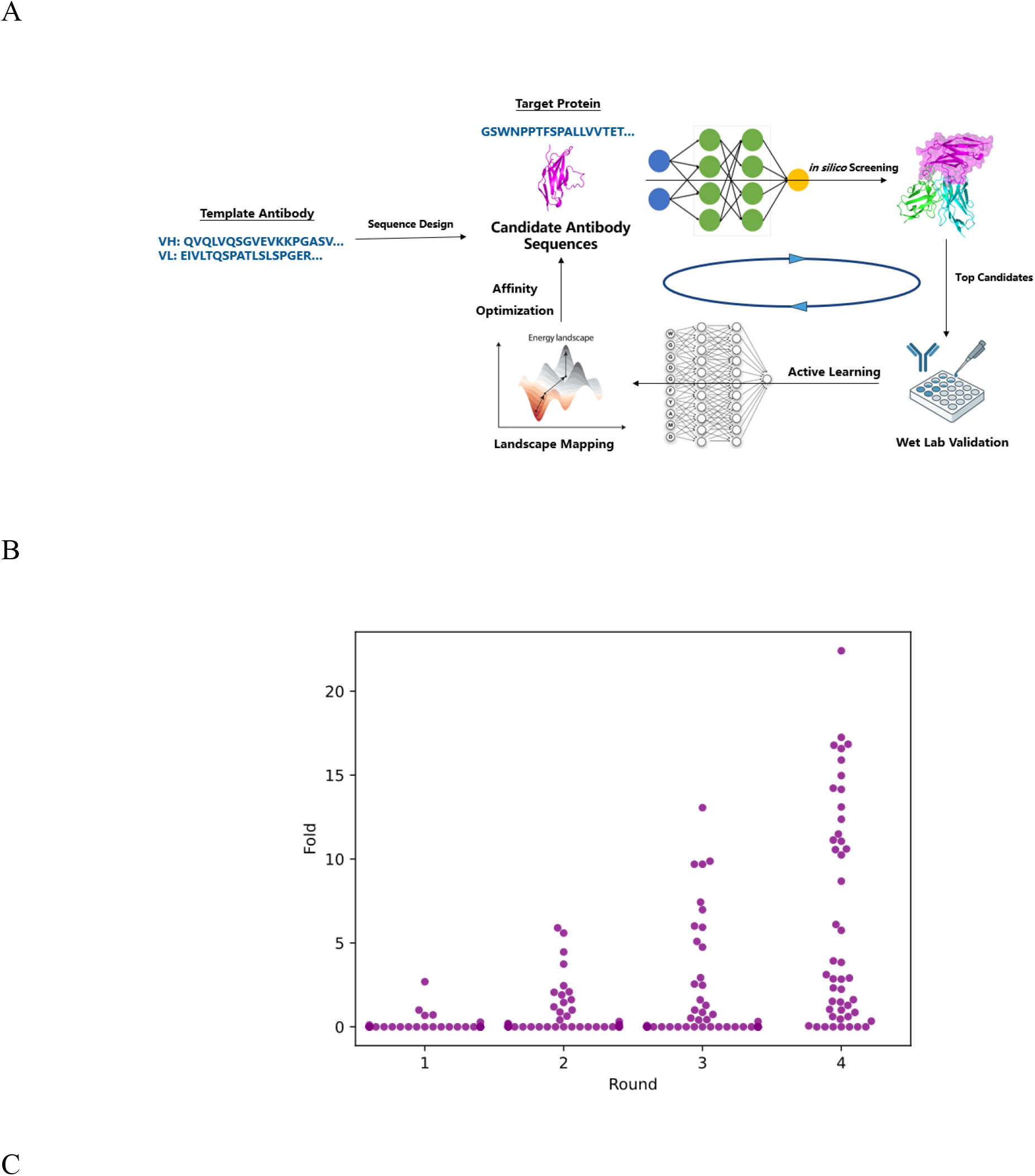
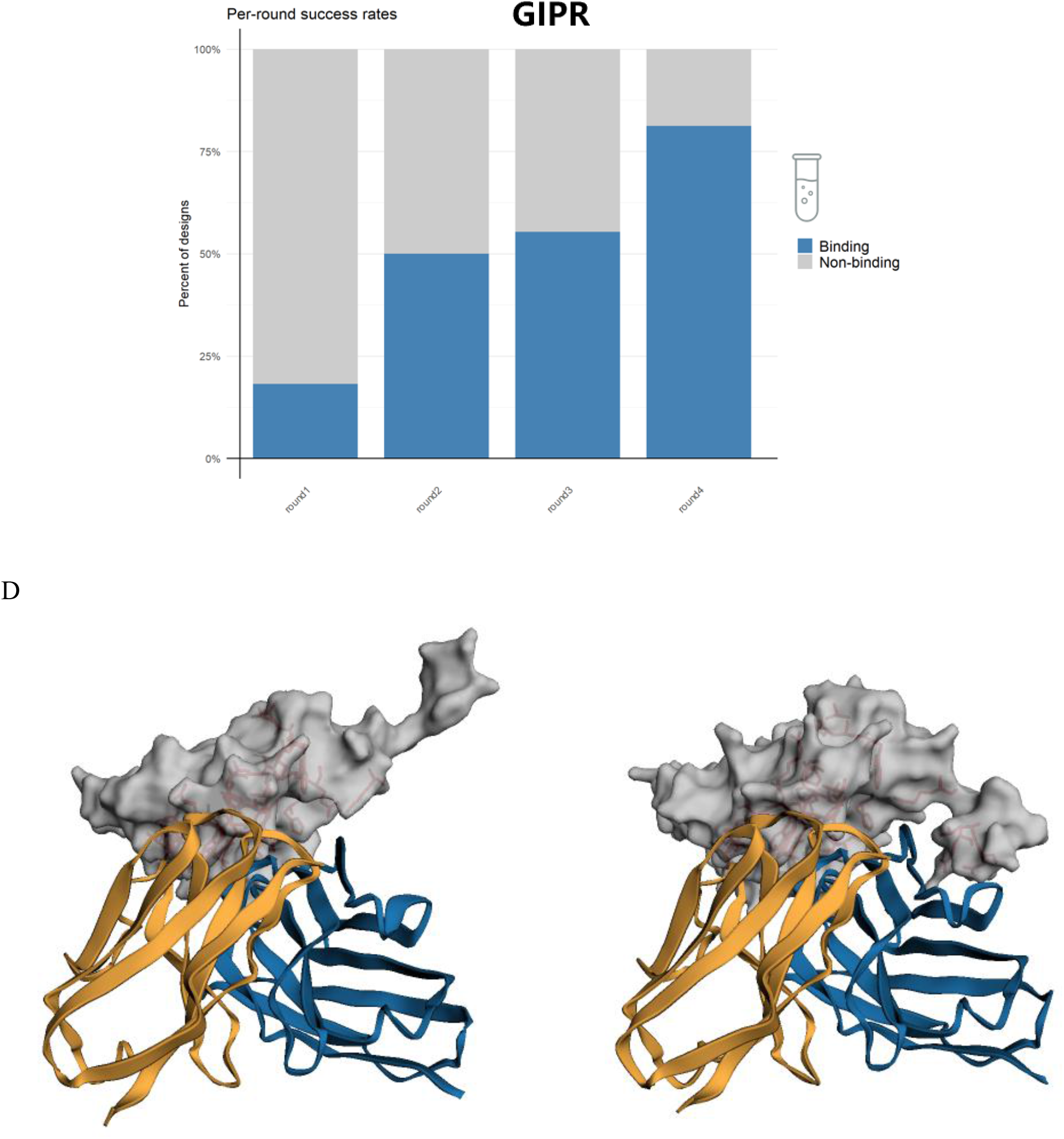
Design and characterization of *in silico* generated GIPR antibodies. (A) Schematic of the iterative active learning process for antibody design. Template antibody sequences and target protein structures were used to generate novel antibody sequences via an iterative active learning approach. Both complementarity-determining regions (CDRs) and framework regions were optimized during the design process. (B) Improvement in binding affinity across design rounds. The ratio of binding affinities of designed antibodies relative to the reference antibody 2G10 is shown for each round. After four rounds of optimization, significant enhancement in binding affinity was achieved. (C) Proportion of designed antibodies with confirmed binding to GIPR across iterative rounds. The percentage of sequences that successfully bind to the target increases progressively from round 1 to round 4, demonstrating the enrichment of functional binders through active learning. (D) Structural comparison of the two lead antibodies, AB106-156 and AB106-131, in complex with GIPR. Left: Predicted binding mode of AB106-156 (heavy chain in blue, light chain in orange). Right: Predicted binding mode of AB106-131 (heavy chain in blue, light chain in orange). GIPR is shown as a gray surface, with key epitope residues highlighted as red sticks. Both antibodies bind to a similar epitope on GIPR, consistent with their comparable functional activity.

By the fourth round, the iterative process produces antibodies with high binding affinities (K_D_ values), and most antibodies showed higher affinity than exceed the benchmark 2G10 in initially screening by Bio-Layer Interferometry (Figure 1B). Notably, the proportion of designed sequences that successfully bind to GIPR increased progressively across rounds, from approximately 30% in round 1 to over 80% in round 4, demonstrating the platform’s ability to enrich for functional binders through iterative learning (Figure 1C). The *in silico* designed antibodies, including AB106-156 and AB106-131, are specifically engineered to bind to a similar epitope on GIPR. Structural predictions reveal that both lead antibodies exhibit overlapping binding modes on the receptor surface (Figure 1D), despite being non-identical to each other or to the reference antibody 2G10 in both the CDR and framework regions (Table 1).

**Table 1.** Sequence identity to 2G10.

| Sample | Sequence Identity | CDR Identity |
| --- | --- | --- |
| AB106-131 | 74.4% | 71.7% |
| AB106-156 | 81.9% | 75.0% |

To ensure favorable drug-like properties, we integrated developability assessment from the outset using our proprietary AlfaDAX platform, which predicts key biophysical and biochemical liabilities including isoelectric point (pI), humanization score (Hu), immunogenicity risk (Immu), aggregation propensity (AP), viscosity risk (Vis), and non-specific binding (NSB). Throughout the iterative design process, we prioritized antibodies with balanced developability profiles. As shown in Supp Table 1, all antibodies selected for experimental validation, including the lead candidates from round 4 (e.g., AB106-131, AB106-156), exhibited low predicted immunogenicity (Immu ≤ 0.1), favorable aggregation and viscosity scores (AP and Vis consistently ≤1.0), and minimal non-specific binding risk (NSB ≤ 0.8). Furthermore, the distribution of these developability metrics across design rounds (Supp Figure 1) demonstrated that the iterative optimization process maintained favorable biophysical properties, with round 4 candidates showing balanced pI values (6–8), high humanization scores (Hu ≥ 0.7), and consistently low risk scores for aggregation, viscosity, and non-specific binding. These *in silico* assessments indicate that our design strategy not only generated high-affinity binders but also ensured high developability, thereby reducing the likelihood of downstream manufacturing and formulation challenges

### 2.2 *In vitro* functional validation of candidate molecules

After four rounds of AI-driven sequence generation and optimization, we obtained sequences completely distinct from the template, with 12 antibody sequences exhibiting substantially improved affinity. These 12 candidates were subsequently evaluated for ELISA and cell binding capabilities using overexpressing cell lines. Leveraging data on *in vitro* molecular activity (Table 2), along with assessments of sequence diversity and developability, two lead candidates, AB106-131 and AB106-156, were ultimately identified. Sequence alignment revealed that AB106-131 shares 74.45% identical amino acids with the variable region of 2G10, while AB106-156 shares 81.94%. Affinity measurements using Biolayer Interferometry (Octet) showed that AB106-131 had an affinity of 1.2 nM and AB106-156 of 1.7 nM (Figure 2A). Under the same experimental conditions, the Affinity of 2G10 was measured as 12 nM, indicating an affinity improvement of approximately 10-fold and 7-fold for the two candidates, respectively. Using ELISA, we compared the binding of the two candidate antibodies and 2G10 to the extracellular domain 1 of GIPR (Figure 2B). HEK293-T cells stably expressing GIPR (P48546) on their surface were used to determine the binding EC_50_ of the antibodies via flow cytometry. The EC_50_ values were 3.2 nM for AB106-156, 2.2 nM for AB106-131, and 2.8 nM for 2G10 (Figure 2C). A cAMP signaling reporter cell line constructed from the overexpressing cell line was used to assess the blockade of the GIP/GIPR signaling pathway by the antibodies. The IC_50_ values for blockade were 5.9 nM for AB106-156, 4.4 nM for AB106-131, and 4.0 nM for 2G10, indicating that the blocking activities of both candidate molecules were comparable to that of 2G10 (Figure 2D). In summary, we have developed novel antibodies that outperform 2G10 in certain aspects.

**Table 2.** Antibody Affinity Measurement Results.

| Sample ID | K<sub>D</sub> (M) | Affinity Ratio | ELISA EC<sub>50</sub> (pM) | Cell Binding EC<sub>50</sub> (nM) |
| --- | --- | --- | --- | --- |
| 2G10 | 1.198E-08 | 1 | 14.14 | 2.36 |
| AB106-122 | 2.424E-09 | 5 | 28.23 | 4.38 |
| AB106-123 | 2.060E-09 | 6 | 70.96 | 4.07 |
| AB106-125 | 1.837E-09 | 7 | 72.91 | 5.64 |
| AB106-131 | 1.196E-09 | 10 | 83.28 | 3.38 |
| AB106-132 | 1.222E-09 | 10 | 48.18 | 3.67 |
| AB106-136 | 2.016E-09 | 6 | 108.3 | 9.59 |
| AB106-144 | 2.308E-09 | 5 | 63.55 | 2.51 |
| AB106-146 | 1.865E-09 | 6 | 68.59 | 4.23 |
| AB106-154 | 1.693E-09 | 7 | 94.22 | 7.82 |
| AB106-155 | 2.449E-09 | 5 | 44.91 | 3.83 |
| AB106-156 | 1.745E-09 | 7 | 93.73 | 2.18 |
| AB106-157 | 2.424E-09 | 5 | 78.59 | 2.15 |

**Figure 2.**
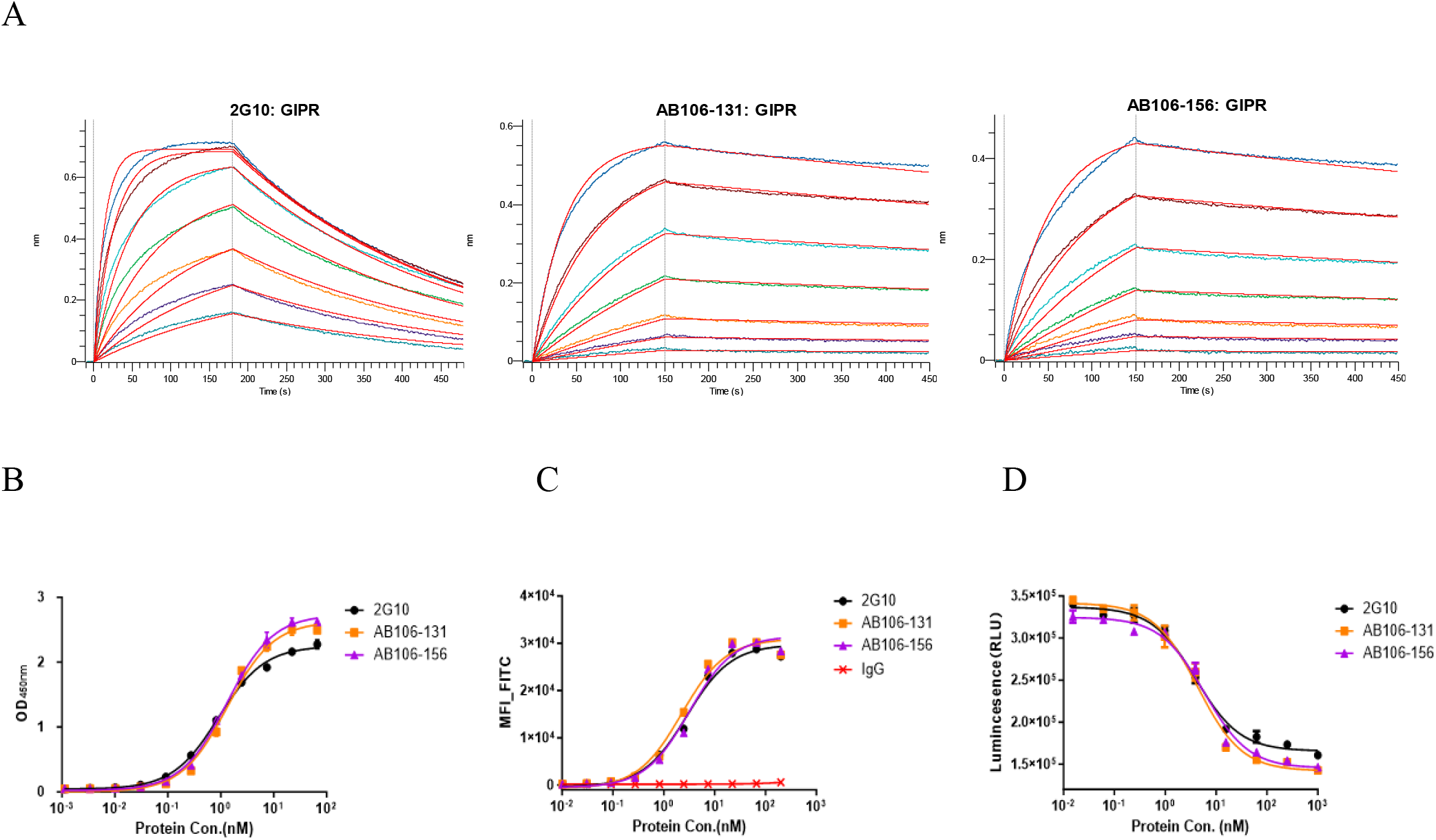
*In vitro* validation data for candidate molecules. (A) Binding kinetics of *in silico* designed antibodies compared to 2G10. Biolayer interferometry (BLI) sensorgrams depict the binding profiles of 2G10 (12 nM) and three designed antibodies AB106-131 (1.2 nM), AB106-156 (1.7 nM). (B) Molecular binding of candidate molecules to the first extracellular domain of GIPR was measured using ELISA. (C) The binding activity of candidate molecules to cells overexpressing GIPR was measured using flow cytometry. (D) The blocking effect of candidate molecules on the cAMP signaling pathway was measured using a human *GIPR* HEK293 luciferase reporter cell line.

### 2.3 *In vivo* efficacy in *hGIPR* DIO mice

To determine the effects of AB106-156 as a monotherapy and in combination with the GLP-1 receptor agonist semaglutide on weight loss, 25-week-old diet-induced obese *hGIPR* mice (DIO) were obtained and maintained on a high fat diet (60 % kcal from fat). Mice received subcutaneous injection of either 0.123 mg/kg semaglutide or equal volume of PBS as vehicle control, 30 mg/kg AB106-156, or a combination of semaglutide and AB106-156 over the course of 25 days (Figure 3A). AB106-156 demonstrated a stronger weight loss effect and data for AB106-131 were therefore omitted and provided in the Appendix. Body weight was monitored twice a week, and body composition was assessed by MRI at days 26 and 44. Over 25 days of treatment, mice treated with semaglutide or AB106-156 lost greater than 15 % of their body weight (Figure 3B), Co-administration of AB106-156 and semaglutide led to -25.4% weight loss, indicating synergistic effect. Body composition analysis revealed an about 36.2 % decrease in fat mass with AB106-156 and a 25.8 % decrease with semaglutide (Figure 3D). Co-administration led to 60.1 % fat mass loss, superior to AB106-156 or semaglutide alone (Figure 3D). Treatment with semaglutide alone decreased lean mass by 5.4% compared to control animals, while AB106-156 alone induced an about 2.2% increase (Figure 3D), indicating treatment of AB106-156 preserved lean mass. However, parameters began rebounding toward baseline at varying rates once treatment ended, Semaglutide rebounded much more bodyweight than AB106-156 (10.7% *vs* 2.7%) by the end of the 19-days washout period (Figure 3B). Body weight rebounded mainly in fat mass following semaglutide discontinuation (Figure 3D), while sustained body weight decrease was observed in mice treated with AB106-156 alone in wash out period. Overall, these data indicate semaglutide alone results in both fat and lean mass loss, while AB106-156 alone results in similar but long-acting weight reduction and fat loss but preserving lean mass.

**Figure 3.**
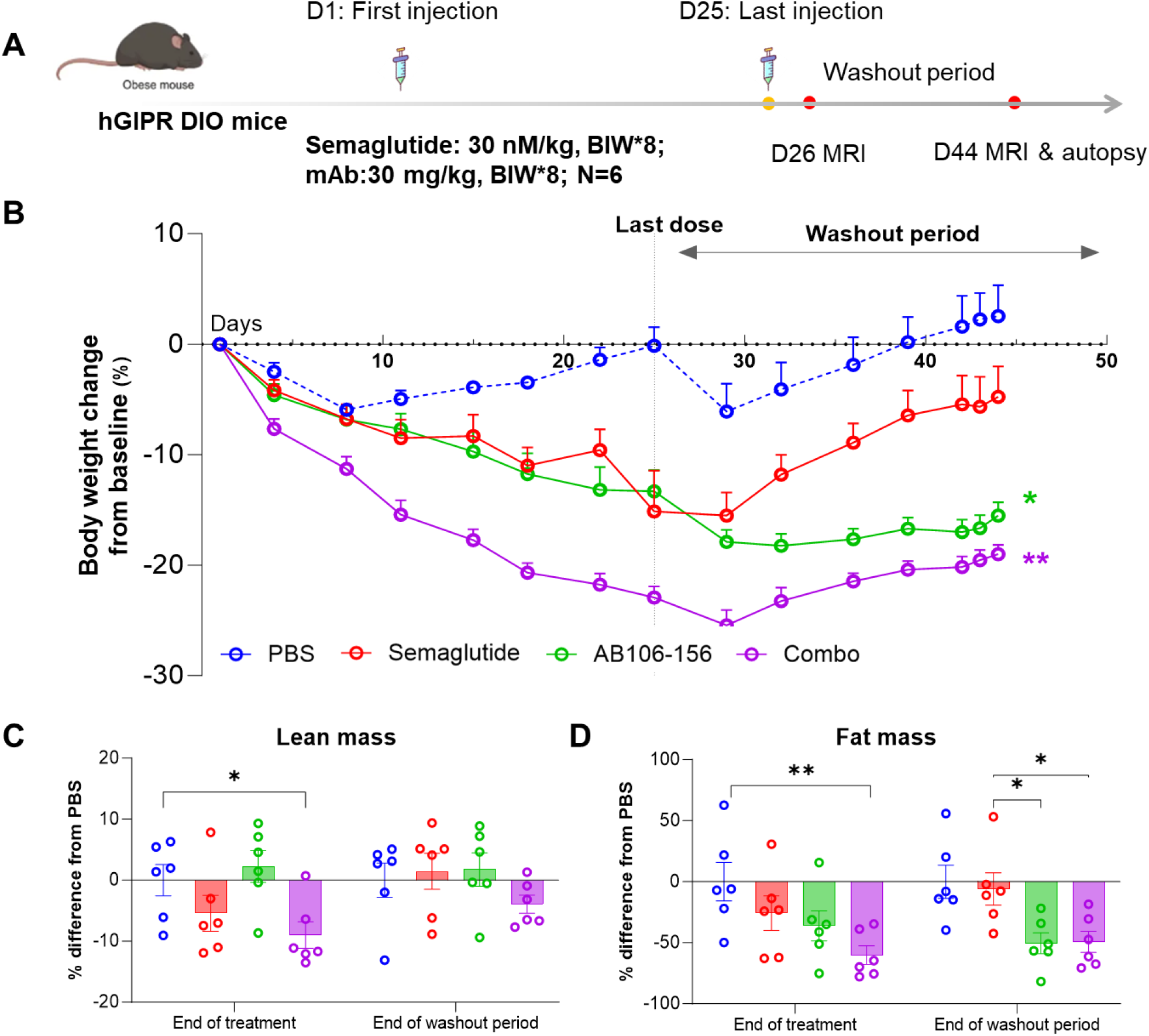
AB106-156 enhanced body weight loss in combination with the GLP-1R agonist semaglutide in DIO mice. (A) Established DIO mice fed HFD for 20 weeks were treated with vehicle, AB106-156 (30 mg/kg, twice a week), and semaglutide (0.123 mg/kg, twice a week) alone or in combination, after 25 days of treatment, all animals were monitored for an additional 19-day washout period (n = 6 per group). (B) Body weight was measured and used to calculate percent change from baseline. (C) Lean mass (D) and fat mass were measured at the end of treatment and washout period, respectively. Data represent means±SEM. Two-way repeated-measures ANOVA both with Sidak’s test for multiple comparisons; \**P* < 0.05, \*\**P* < 0.01.

At the end of the study, adipose tissue depots were isolated and weighed. Consistent with the body weight results, the AB106-156 treatment group showed significant reductions in inguinal and epididymal fat pads, with retroperitoneal and brown adipose tissues also reduced by 30% to 50%, respectively. Similar effects were observed in the combination therapy group, indicating that the antibody played a major role. Metabolic parameters were also measured, and all treatment groups showed decreasing trends in low-density lipoprotein cholesterol (LDL) and total cholesterol (TC). Interestingly, the GIPR antibody AB106-156 group exhibited a smaller rebound in fasting insulin levels, indicating improved insulin sensitivity. At the end of the treatment phase and washout period, animals also underwent an Oral Glucose Tolerance Test (OGTT). Compared with the PBS control group, the area under the curve (AUC) of blood glucose was significantly reduced in the combination therapy group, indicating that AB106-156 did not worsen glucose tolerance (data not shown).

## 3 Methods

### 3.1 Protein Expression

The AI-recommended amino acid sequence was optimized for the codon preference of *Cricetulus griseus* (CHO). The heavy and light chains of the antibody were separately constructed into the pCDNA3.4 vector through gene synthesis. The heavy and light chain plasmids were co-transfected into CHO-K1 cells for cultivation and expression. The supernatant of the expressed cells was harvested and purified using ProteinA (Cytiva,) purification resin.

### 3.2 Affinity determination of candidate molecules

Antibody-antigen binding kinetics were assessed by Bio-Layer Interferometry using an Octet® (R8) instrument (Sartorius). All steps were performed at 30°C with shaking at 1,000 rpm in kinetics buffer [PBS, pH 7.4, 0.02% Tween-20]. Antibodies were captured onto ProA biosensors (18-5010, Sartorius) to a level of 4.0 nm. The loaded sensors were then exposed to a concentration series of the antigen (ranging from 0 to 75 nM) for 150 seconds to measure association (Ka), followed by a 300-second dissociation step in buffer (Kdis). Sensorgrams were reference-subtracted and globally fitted to a 1:1 binding model using the instrument’s proprietary software to calculate the affinity constant (K_D_).

#### 3.3 ELISA detects antibody binding to GIPR

The binding of in-house purified antibody to GIPR was assessed by ELISA. Recombinant GIPR antigen (18774-H08H, Sino Biological) was coated onto 96-well plates at 0.5 μg/mL overnight at 4°C. After blocking with BSA, serially diluted antibody was added and incubated. Bound antibody was detected using HRP-conjugated Mouse Anti-Human IgG Fc secondary antibody (Cat. No. A01854, GenScript) followed by TMB substrate; absorbance was read at 450 nm. EC_50_ values were calculated by four-parameter logistic curve fitting using GraphPad Prism.

#### 3.4 Flow cytometry assay of hGIPR antibody binding activity

The binding of the antibody to GIPR was evaluated by flow cytometry using *GIPR*-overexpressing HEK 293 cells. Cells were harvested and washed with PBS containing 1% BSA-PBS. Approximately 1×10 ^6^ cells per sample were incubated with serial dilutions of the antibody in staining buffer for 60 minutes at 4°C. After washing twice with staining buffer, cells were incubated with FITC-conjugated Goat Anti-Human IgG Fc secondary antibody (Cat. No. ab97003, Abcam) for 60 minutes at 4°C in the dark. Cells were washed again and resuspended in PBS for analysis. Fluorescence intensity was measured using a FongCyte flow cytometer (ChallenBio, China). Data was analyzed using the device’s analysis software, and the mean fluorescence intensity (MFI) was plotted against antibody concentration. EC_50_ values were calculated by four-parameter logistic curve fitting using GraphPad Prism.

### 3.5 hGIPR antibody functional activity by cAMP assay

The inhibitory effect of the antibody on cAMP signaling was assessed using a luciferase reporter gene assay in *GIPR*-overexpressing HEK 293 cells stably transfected with a cAMP-responsive element (CRE)-driven luciferase reporter. Cells were cultured in complete medium, harvested, and resuspended in assay buffer (e.g., PBS containing 0.1% BSA). Cells were seeded into white 96-well plates at a density of approximately 7 × 10^4^ cells per well in serum-free medium. Serially diluted antibody was added to the wells and pre-incubated for 60 minutes at 37°C. To stimulate cAMP production, a sub-maximal concentration of GIPR agonist (GIP peptide, GenScript) was added to each well at a predetermined EC_80_ concentration (0.05 nM), and the plate was incubated for an additional 6 hours at 37°C in a CO_2_ incubator. Following incubation, luciferase activity was detected by adding an equal volume of luciferase assay reagent (GM-040505C, GMBrightOne-Step, Genomeditech) to each well. Luminescence was measured using a multimode microplate reader (Infinite 200 pro m plex, Tecan). Data was analyzed using GraphPad Prism software. Luminescence intensity (relative light units, RLU) was plotted against antibody concentration, and the half-maximal inhibitory concentration (IC_50_) was calculated by four-parameter logistic curve fitting.

### 3.6 Animal study

#### Test article

Semaglutide used in the study was Ozempic (Novo Nordisk, 202409CCE1), and the monoclonal antibody was manufactured by Gene Universal (Anhui) Co., Ltd.

#### Diet and grouping

Male *hGIPR* mice aged 6–7 weeks old (Cat#112714) was purchased from Biocytogen and the study was performed according to the regulations and SOP of Biocytogen. Animals were group housed and maintained on a 12 h light/12 h dark cycle at standard temperature and humidity conditions and fed a high-fat diet *ad libitum* for 15 weeks prior to the start of study.

#### Pharmacodynamic assay

Mice were randomized into four groups based on body weight and body fat percentage and were subcutaneously administered one of the following treatments: Negative control (NC, commercially sourced PBS solution); AB106-156 (twice a week); Semaglutide (0.123 mg/kg, twice a week), combo treatment (AB106-156 & Semaglutide), respectively. Body composition was assessed using a Low-field NMR body composition analyzer (*Niumag Analytical, Suzhou*), measured at the end of last injection and the end of wash out period. Cumulative food intake was determined by manually weighing food from single cages twice a week. Body weight was measured twice a week. Mice were euthanized on days 44 of the study.

### 3.7 Statistical analysis

One- or two-way ANOVA followed by Sidak’s test for multiple comparisons (with repeated measures for time series data) was used in all studies. For comparison among multiple groups at a single time point, one-way ANOVA was performed. All tests used the software GraphPad Prism. Significance was defined as \**P* < 0.05, \*\**P* < 0.01, and \*\*\**P* < 0.001.

## 4 Discussion

GIPR and GLP-1R have been firmly established as clinically relevant targets for obesity management, as evidenced by the regulatory approval and widespread clinical adoptions. While a growing pipeline of multi-target therapeutics targeting GIPR, GLP-1R, and other GPCRs is currently under investigation, mostly are peptide-based characterized by short half-life. The approved injectable anti-obesity agents-once-weekly tirzepatide (Zepbound®), semaglutide (Wegovy®), and daily liraglutide (Saxenda®), yield weight reductions spanning 6% to 20%^[23]^. This efficacy hierarchy underscores a paradigm shift toward less frequent dosing and multi-receptor targeting in obesity pharmacotherapy. As for individuals with obesity, less frequent dosing and greater ease of administration are key factors in improving treatment adherence and long-term outcomes.

By contrast, the conjugation of GIPR antibody with GLP-1R agonists or other peptide moieties represents a promising strategy to markedly prolong systemic half-life, thereby enabling extended dosing intervals. AMG133, a bispecific molecule engineered by conjugating a fully human monoclonal anti-human GIPR antagonist antibody to two GLP-1 analogue agonist peptides using amino acid linkers, represents a novel therapeutic approach distinguished by its extended pharmacokinetic profile. It’s administered on a once-monthly schedule with a half-life of approximately 21 days^[24]^, which is three times that of currently marketed once-weekly agents (tirzepatide and semaglutide). Preliminary clinical findings suggest that weight loss achieved with AMG 133 is comparable to that of tirzepatide and exceeds that of GLP-1R monoagonists. Notably, the absence of a plateau in weight reduction at 52 weeks indicates that the full therapeutic potential has yet to be realized. Collectively, these attributes support the possibility of further extending the dosing interval, with future investigations exploring maintenance dosing every two months or longer after target weight loss is attained. The sustained weight reduction without plateau may be attributed to the extended half-life of the antibody, which enables prolonged target engagement with less frequent dosing.

In this study, we employed an iterative active learning platform, AlfaBodY, to *in silico* design human antibodies targeting GIPR. By integrating structural and sequence information, we generated antibodies with high binding affinities (K_D_ values as low as 1.2 nM). Two lead candidates, AB106-131 and AB106-156, exhibited approximately 10-fold and 7-fold higher affinity than 2G10, respectively, while maintaining comparable antagonistic activity in *GIPR*-overexpressing HEK 293 cells by cAMP reporter assay (IC_50_ 4∼5 nM), similar to that of the 2G10.

The *in vivo* efficacy of AB106-156 is particularly noteworthy. In diet-induced obese (DIO) mice, AB106-156 achieved a maximum weight reduction similar to that of semaglutide, indicating comparable efficacy between the two agents after the last treatment. The 25-day treatment with AB106-156 alone led to >15% weight loss in DIO mice, an effect that was amplified by co-treatment with semaglutide, resulting in a 25.4% reduction. This result differs from published literature, in which GIPR antibody generally shows minimal weight loss, highlighting the unique pharmacological profile of the antibody generated in this study. Surprisingly, the weight loss achieved by AB106-156 was primarily driven by a reduction in fat mass, with a parallel increase in lean mass, thereby contrasting sharply with the loss of lean mass observed following semaglutide monotherapy. Regarding durability and the prevention of weight rebound, AB106-156 showed significant advantages over peptide-based therapies. Following a 19-day washout period, a marked divergence in durability was observed between the GIPR antibody and semaglutide. The semaglutide group experienced a substantial weight gain (rebounding from -15% to -4%), representing a 10.7% increase. This was accompanied by a substantial increase in fat mass, a finding consistent with the short half-life inherent to peptide-based agents. In contrast, AB106-156 monotherapy maintained a 15% weight reduction, preserving most of its maximal weight loss (-18%), showed minimal weight rebound (+2.7%). The combination group likewise exhibited robust durable weight loss, a finding that may be largely attributable to the prolonged half-life of the antibody. These findings align with the clinical observations for AMG133, a bispecific molecule combining GIPR antagonism with GLP-1 receptor agonism, which demonstrated up to 20% weight loss after 52 weeks and a durable effect with monthly dosing^[25]^. Our results suggest that a pure GIPR antagonist, when combined with a GLP-1 agonist, can recapitulate the synergistic efficacy while potentially offering flexibility in dosing regimens. The sustained effect after washout may reflect a unique physiological consequence of GIPR blockade, such as alterations in adipose tissue homeostasis or central appetite regulation, which merit further investigation. These results suggested the long-acting efficacy of AB106-156 supports less frequent dosing when compared with peptide-based agents that require frequent administration, representing a significant advantage in real-world clinical settings.

Our results compare favorably with recent advances in computational design of GPCR-targeting biologics. Baker and colleagues recently described the *de novo* design of miniprotein antagonists for several class B GPCRs, including GIPR, using RFdiffusion and metaproteome-derived scaffolds^[26]^. Through high-throughput yeast display screening of thousands of designs, they identified miniproteins with nanomolar to picomolar affinities for the GIPR extracellular domain and confirmed their antagonistic activity in cAMP assays. Antibodies generated through *de novo* approaches often have unknown *in vivo* efficacy and potential risks. In addition, such high-throughput screening typically requires certain display platforms and library synthesis capabilities, and the one-to-one pairing and expression of a large number of antibody heavy and light chains is technically challenging. In contrast, our approach starts from clinically validated antibodies to further optimize and achieve better efficacy, and offer potential advantages in clinical development, including reduced immunogenicity and increased developability. Furthermore, whereas the Baker study focused on binder discovery and *in vitro* characterization, we have extended our evaluation to a preclinical efficacy model, demonstrating that an *in silico* designed anti-GIPR antibody can achieve significant weight loss and favourable metabolic effects *in vivo*.

Despite the substantial improvement in binding affinity relative to 2G10, the functional antagonism of our GIPR antibodies (IC_50_ for cAMP blockade) was not proportionally enhanced; it remained comparable to the reference. This observation underscores a recurring challenge in developing functional antibodies against GIPR: high-affinity binding does not always translate into improved antagonistic potency, possibly due to the complex conformational dynamics of the receptor or the necessity of precisely occluding the orthosteric peptide-binding site. Notably, using the same AlfaBodY platform we have successfully enhanced the functional activity (5-to 10-fold) of antibodies against other targets, such as MSTN, BTLA, and ActRII (unpublished data), indicating that the platform’s capability to optimize function is target-dependent. For GIPR, future designs may require focusing on epitopes that more effectively disrupt receptor activation, perhaps by targeting both the extracellular domain and transmembrane segments, or by engineering antibodies that lock the receptor in an inactive conformation.

Several limitations of this study should be acknowledged. First, the *in vivo* efficacy was evaluated in a humanized *GIPR* mouse model; extrapolation to humans requires caution. Further, the effective dose and dose-response relationship are not clear for the time being, a multi-dose pharmacodynamic study will be initiated. Second, the combination therapy employed separate administration of antibody and semaglutide, the dosage of AB106-156 and semaglutide selected in the study were widely used in similar experiments to ensure maximal target coverage and weight loss effects and maybe higher than that initial pharmacodynamic study. Also, a bispecific format like AMG133 might offer practical advantages for clinical translation, we cannot conclude that similar weight loss effect was achieved under equimolar dose. Third, while our antibodies demonstrated favourable developability profiles *in silico* (AlfaDAX scores), additional experimental assessments of immunogenicity, viscosity, and long-term stability are needed.

In conclusion, we have successfully applied an AI-driven active learning platform to design novel anti-GIPR antibodies with high affinity and potent *in vivo* efficacy. The lead antibody, AB106-156, alone or in combination with a GLP-1R agonist, achieved sustained weight loss without rebound in DIO mice compared to the GLP-1R agonist alone, highlighting the therapeutic promise of GIPR antagonism. Our work also illustrates that while high-affinity binding can be readily achieved, functional optimization remains target-specific and may require tailored design strategies. Future efforts will focus on engineering bispecific molecules that integrate GIPR antagonism with GLP-1R agonism, developing ultra-long-acting, non-rebound weight loss bifunctional molecules, and on advancing lead candidates toward clinical evaluation for obesity and related metabolic disorders.

## Supplementary

Supp Table 1. DAX scores of designed antibodies of 4 Round.

**Supp Figure 1.**
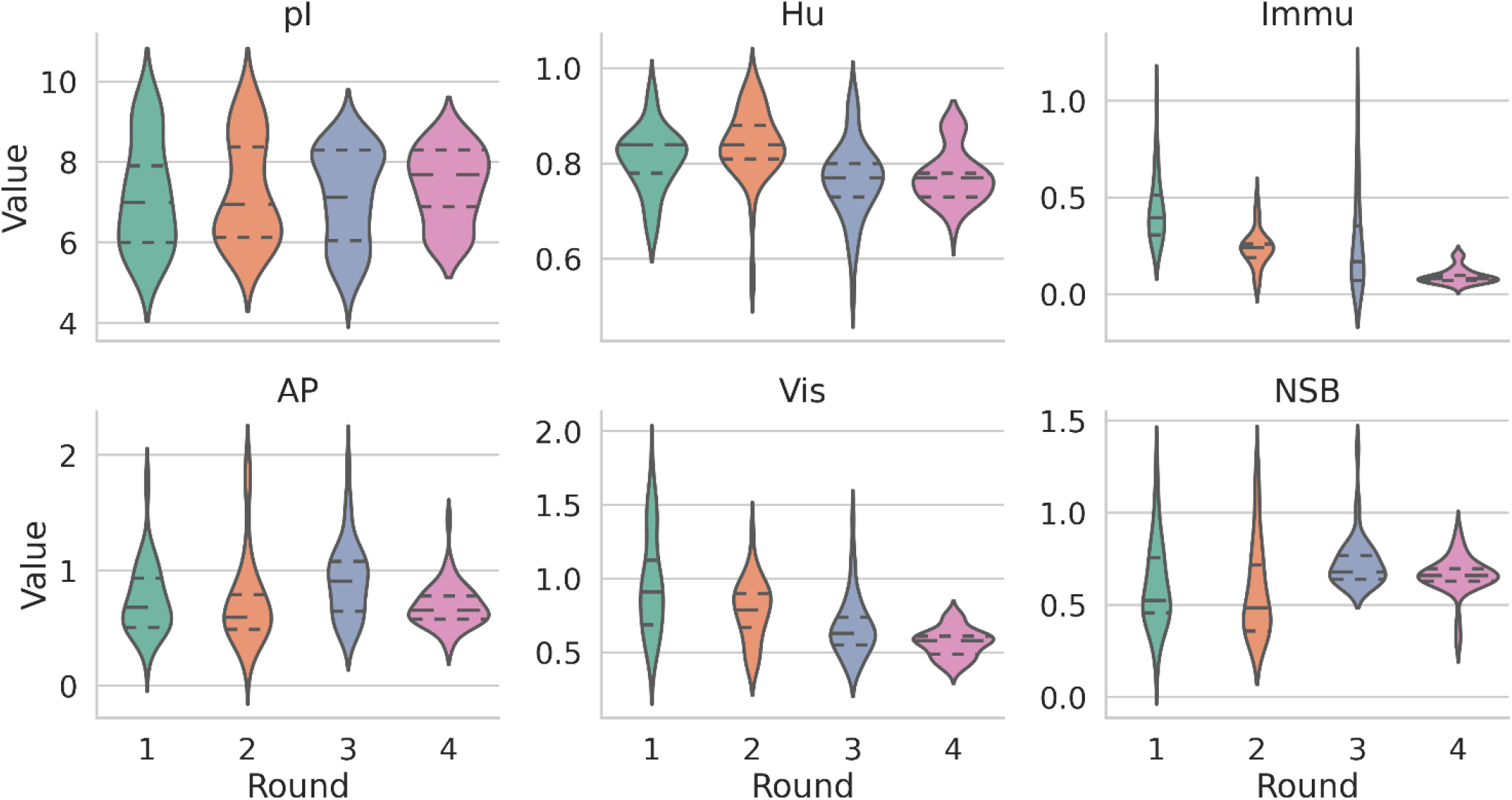
*In silico* developability metrics of designed antibodies across iterative design rounds. Violin plots showing the distribution of key developability parameters for antibodies generated in rounds 1 to 4, as predicted by the AlfaDAX platform. Parameters include: **(A)** Isoelectric point (pI); **(B)** Humanization score (Hu); **(C)** Immunogenicity risk (Immu); **(D)** Aggregation propensity (AP); **(E)** Viscosity risk (Vis); **(F)** Non-specific binding (NSB). The central horizontal line represents the median, and the dashed lines indicate the quartiles. For Immu, AP, Vis, and NSB, scores ≤ 1 are considered low risk. Across all rounds, round 4 antibodies maintained favorable developability profiles, with pI values within the typical therapeutic antibody range (6–8), high humanization scores (Hu ≥ 0.7), and all risk scores consistently below the threshold of 1.0, indicating low liabilities for immunogenicity, aggregation, viscosity, and non-specific binding.

**Supp Figure 1.**
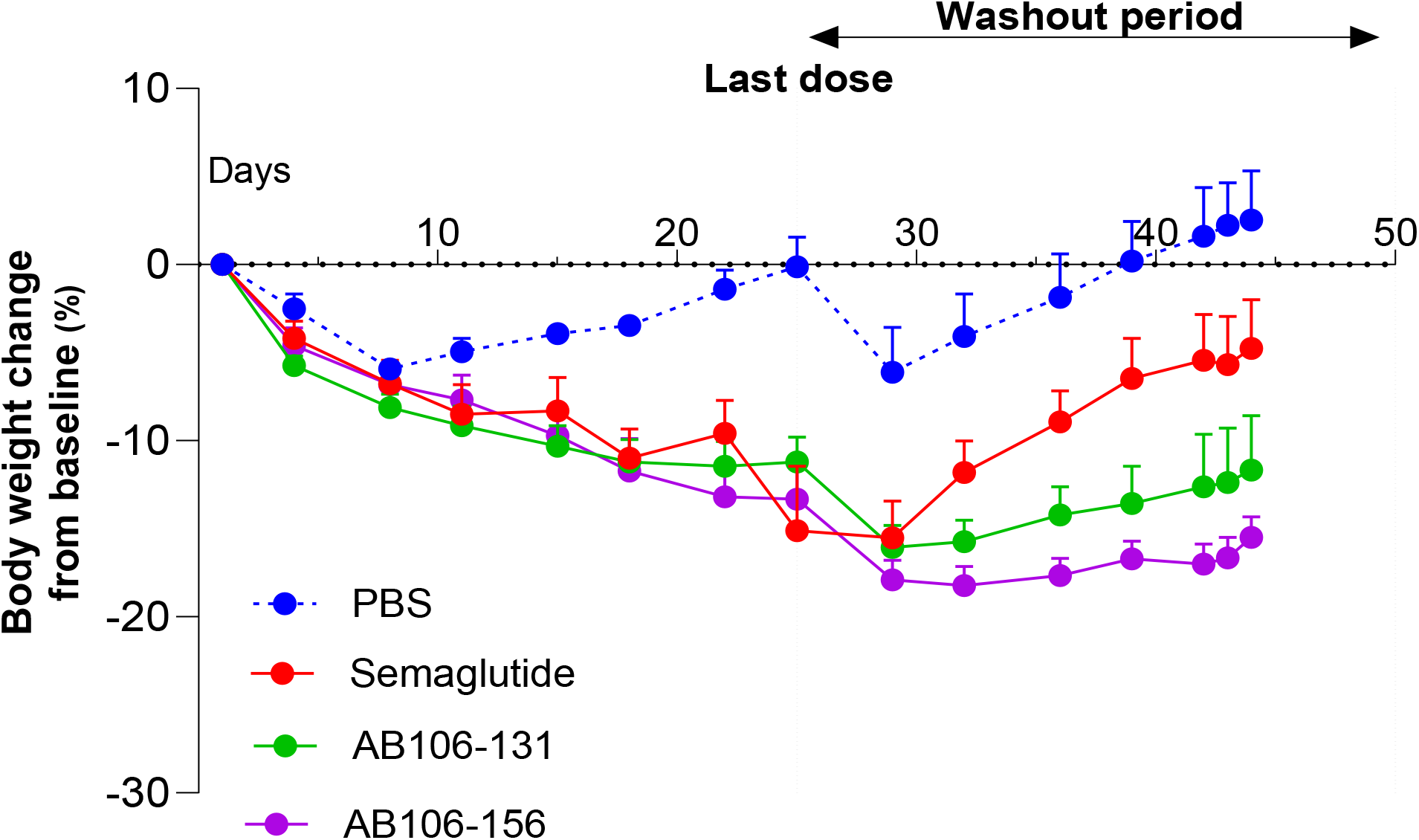
Efficacy results for AB106-131. Comparison of body weight reduction between AB106-131 and AB106-156 in diet-induced obesity (DIO) mice, and AB106-156 exhibited superior weight reduction.

